# Reward Prediction Error Signaling during Reinforcement Learning in Social Anxiety Disorder is altered by Social Observation

**DOI:** 10.1101/821512

**Authors:** Michael P. I. Becker, Rolf Voegler, Jutta Peterburs, David Hofmann, Christian Bellebaum, Thomas Straube

## Abstract

**BACKGROUND:** Dysfunctional expectations of impending social or performance outcomes are core features of Social Anxiety Disorder (SAD) but often lack formal definition in clinical research. Reinforcement learning (RL) models offer a framework to define changes in outcome expectations in a formal way by computing the prediction error (PE). This study quantifies the updating of expectations by PEs in SAD and investigates alterations in RL regions associated with PE signaling.

**METHOD:** 48 adult participants (24 diagnosed with SAD and 24 age-, gender-, and education-matched healthy controls (HC)) underwent event-related functional magnetic resonance imaging while learning from probabilistic feedback. Crucially, both groups completed two parallel versions of the task: one in which they learned under scrutiny (social observation) and one in which they learned without being overtly evaluated (non-social control condition).

**RESULTS:** Coupling to prediction errors in SAD was elevated in dorsomedial prefrontal cortex (DMPFC) when learning under observation. These findings provide the first evidence that PE signaling during social performance situations in SAD is associated with hypersensitive response signatures in DMPFC, a brain region associated with using others’ value standards as a proxy for one’s own value standards. Dynamic Causal Modelling further revealed that RPE-modulated connectivity from ventral pallidum to DMPFC during observation was reduced in SAD.

**CONCLUSIONS:** The present results corroborate a crucial role of DMPFC in SAD, which corresponds to dysfunctional expectations about others’ alleged performance standards that play a prominent role in current models of the disorder.

## Introduction

Expectations of failure and persistent fear responses during social performance or interactions characterize Social Anxiety Disorder (SAD; American Psychiatric Association, 2013). Patients suffering from SAD most often dread exposure to scrutiny by others, set exaggerated standards for the success of their own social behavior, and invest a lot of time *ex post facto* to recapitulate negative aspects of past interactions (Rapee & Heimberg, 1997). Functional models of social anxiety and its clinical manifestation, SAD, converge on the maintaining role of dysfunctional expectations in the development and exacerbation of avoidance and anxiety (Clark & Wells, 1995; Rapee & Heimberg, 1997). In particular, dysfunctional expectations about impending social or performance situations are core features of SAD. In general, altered expectations can result from avoidance behaviors and help reduce anxiety (Lovibond et al., 2008). In SAD, situation-specific expectations bias predictions regarding the evaluation of one’s actions by others in social situations (Rief et al., 2015, Rapee & Heimberg, 1997). Hence, negative predictions of one’s own performance in social situations supposedly increase the risk of SAD-related avoidance behavior and might even foster maladaptive social strategies that increase the risk of negative evaluation (see reviews by Rief et al., 2015, Hofmann, 2007, Clark & Wells, 1995). The degree of alteration in these predictions might further affect the course and severity of the disorder as well as responsiveness to therapy (Safren et al., 1997; Delsignore & Schnyder, 2007; El Alaoui et al., 2015).

The updating of predictions is a mechanism that is hypothesized to be altered in anxiety disorders (see reviews by Grupe & Nitschke, 2013, Cavanagh & Shackman, 2015, Paulus & Stein, 2006, Paulus & Yu, 2012). In particular, altered updating may be characterized by (a) altered prediction errors (White et al., 2017, Homan et al., 2019) and (b) altered learning rates for disorder-relevant stimuli (Browning et al., 2015, Koban et al. 2018, Homan et al., 2019) as formalized by reinforcement learning (RL) models. Specifically in SAD, we would expect excessive prediction errors to performance feedback as well as lower learning rates if the updating of predictions takes place in socially evaluative contexts. Previous research in subclinical anxiety, Post-traumatic Stress Disorder, and SAD has shown that individual differences in learning rates predict individual differences in symptom severity (Browning et al., 2015, Homan et al., 2019, Koban et al. 2017).

RL allows for the investigation of prediction errors on the trial level as well as of overall learning rates but requires a framework of probabilistic learning to do so. Probabilistic learning in SAD has previously been investigated behaviorally (Koban et al. 2017) as well as with respect to its electrophysiological correlates (Voegler et al., 2019). Voegler et al. (2019) found that SAD patients learned better from negative feedback in a control condition and learned worse from negative feedback in a socially evaluative context. In addition, Koban et al. (2017) found that SAD patients did not exhibit the positivity bias present in control subjects and that patients’ fear of negative evaluation was negatively associated with learning rates. Moreover, a number of studies have investigated subclinical social anxiety (Abraham & Herrmann, 2015, Stevens et al., 2014, Piray et al., 2019, Hunter et al., 2019). These studies generally report altered learning with regard to socially evaluative information while presenting mixed findings regarding altered learning rates in individuals with heightened social anxiety. However, subclinical symptom severity is relatively low in these studies and their relevance for clinical manifestation of SAD remains questionable. In sum, a growing body of evidence suggests that feedback learning and updating are biased towards negative information in clinical and non-clinical social anxiety.

Reward Prediction Errors (RPEs) have been shown to be associated with activation of a reward-learning network (RLN) comprising ventral striatum (VS), lentiform nucleus, ventromedial prefrontal cortex (VMPFC), and midbrain nuclei (Haber & Behrens, 2015; Pauli et al., 2018). Additionally, RPEs coding for social variables like others’ advice (Behrens et al., 2008) or other’s value standards (Nicolle et al., 2012) are coupled to activation of a network associated with mentalizing of others’ goals and intentions (, Wittmann et al., 2016; Schurz et al., 2014), comprising Area 9 of dorsomedial prefrontal cortex (DMPFC) and temporoparietal junction. DMPFC has also been shown to respond to the perception of scrutiny (Gimenez et al., 2012, Becker et al., 2014). In SAD, socially evaluative contexts trigger differential responses in VS (Becker, Simon, et al., 2017) as well as in regions of medial prefrontal cortex (MPFC, Pujol et al., 2013), and alter functional connectivity patterns (Gimenez et al., 2012, Heitmann et al., 2016). In particular, performance feedback given to SAD patients during social observation (Becker et al., 2017) has been linked to these alterations in network activation. Similar findings from healthy subjects with heightened but subclinical social anxiety suggest that MPFC activation is blunted during learning and feedback processing (Peterburs et al., 2016; Piray et al., 2019). However, it is as yet unknown if patients with clinically significant SAD show alterations of RL network activation during prediction error signaling and updating. More importantly, we do not know if RPE-related network activation has a role in triggering biased expectations regarding the evaluation by others in social situations.

A critical aspect of a socially evaluative situation is the immanent feeling of being observed by someone. In order to trigger this feeling, we used an observation manipulation during a probabilistic feedback-learning task (Frank, 2004) to investigate if indices of updating during RL are altered in socially evaluative contexts. Specifically, we expected that updating of learned associations takes place at a slower rate during observation in SAD. Behaviorally, lower learning rates in SAD point towards slower updating in more anxious individuals (Browning et al., 2015). The probabilistic feedback-learning task also allows assessing the differential contributions of positive feedback and negative feedback on learning. Based on our previous research (Voegler et al., 2019), we hypothesized that HC would learn better from positive feedback, while SAD patients would show better learning from negative feedback, especially in the observation condition. On the brain functional level, RPEs during feedback presentation were expected to predict activation of a reward-learning network comprising ventral striatum (VS), lentiform nucleus, and ventromedial prefrontal cortex (VMPFC) (Haber & Behrens, 2015; Pauli et al., 2018). We expected observation to modulate RPEs in SAD patients. Analogous to the socio-emotional hypersensitivity they typically exhibit in evaluative social situations and based on findings from several lines of research, we expected SAD patients to show hypersensitive response signatures in medial prefrontal cortex (Voegeler et a., 2019; Bas-Hogendam et al., 2019, Heitmann et al., 2016). Based on these studies’ coordinates, we hypothesized an interaction of observation and PEs would recruit Area 9 of DMPFC. We also assumed that activation in VS and lentiform nucleus would show this group-by-observation interaction. Recent studies have shown differences between SAD and HC in VS activation to different feedbacks implying that these differences might indicate blunted signaling of positive Pes (Boehme et al., 2014, Becker et al., 2017). Other studies have shown differences between generalized anxiety disorder and HC in activation of lentiform nucleus, implying that these differences might indicate excessive signaling of negative PEs (White et al., 2017).

As RPEs are a likely candidate mechanism for the modulation of connection strength between regions (den Ouden et al., 2011), another aim of the present study was to assess differences in effective connectivity between SAD and HC. In an earlier study, Heitmann et al. (2016) identified aberrant functional connectivity in both DMPFC and VS in SAD. Based on their findings, we used Dynamic Causal Modelling to model dependencies between the mentalizing network and the reward network. A hierarchical analysis based on Parametric Empirical Bayes (PEB; Friston et al., 2016) allowed us to model and test the following predictions while controlling for between- and within-subject variances: (1) the RPE modulates the effective connectivity between the mentalizing network and the reward network, (2) observation gates this connectivity, and (3) this gating is amplified in SAD. Specifically, we tested a model containing RPEs as modulatory input against a null model that only featured valence (thus disregarding the magnitude information of the RPE).

## Methods and Materials

### Sample characteristics

Participants were recruited at the Institute of Medical Psychology and Systems Neuroscience at the University of Muenster, Germany. fMRI data from 24 patients with a DSM-IV diagnosis of SAD (mean age: 25.42 ± 2.78 years; 14 females) and 24 healthy control subjects (HC) without any diagnosis (mean age: 24.46 ± 4.17 years; 16 females) were analyzed. HC were matched to SAD patients with regard to age, gender, and years of education. Before participation, all subjects were screened for neurological and mental disorders. Exclusion criteria for both groups included a history of or currently present psychotic, substance-related, or neurological disorders, in particular severe medical conditions that might influence neurocognitive function (e.g., head injury with loss of consciousness).

The German version of the Structured Clinical Interview (SCID) for DSM-IV (SKID-I; Wittchen et al., 1997) was used to assess patients’ current diagnostic status. All interviews were conducted by an experienced clinical psychologist (R.V.). None of the subjects fulfilled the criteria for a diagnosis of a current episode of major depression, obsessive-compulsive disorder, general anxiety disorder, eating disorder, psychotic disorder, or substance abuse as defined by the SCID. Four SAD patients reported previous episodes of major depressive disorder. Six of the SAD patients were in cognitive-behavioral treatment and two received psychopharmacological treatment (both with citalopram). HC and SAD were administered the Beck-Depression Inventory (BDI-II, Beck, et al., 2006), the Social Phobia Scale (SPS), the Social Interaction Anxiety Scale (SIAS, Mattick and Clarke, 1998), and the self-report version of the Liebowitz Social Anxiety Scale (LSAS, Liebowitz, 1987).

Written informed consent was obtained before starting the experimental procedure. The study procedure conforms to the ethical standards of the Declaration of Helsinki and was approved by the Ethics Committee of the German Psychological Society (Deutsche Gesellschaft für Psychologie, DGPs). All subjects received monetary reimbursement for participation, including additional incentives based on learning performance in the experimental task (see next section). All participants, except one, also participated in a separate EEG study (Voegler et al., 2019).

### Experimental design

#### Observation manipulation

Before starting the experimental procedure, controls and patients were informed that they would be exposed to two different experimental conditions in two separate task runs. In one condition they would be observed with a video camera that was mounted to the head coil (observation condition), with the video feed being watched by an observer who would critically evaluate their performance. In the other condition, the camera was not mounted and subjects were explicitly informed that there would be no observation (control condition).

After each run, participants were asked to rate their arousal level and feelings of unpleasantness related to positive and negative feedback on Likert scales from 1 (low arousal/not unpleasant at all) to 9 (extreme arousal/extremely unpleasant). After the observational run, this questionnaire included a question that assessed how uncomfortable subjects had felt while being observed, also on a Likert scale ranging from 1 (not uncomfortable at all) to 9 (extremely uncomfortable).

#### Probabilistic learning task

The experimental paradigm was an adapted version of the probabilistic learning task described by Frank et al. (2004). Participants learned to associate pairs of Japanese Hiragana characters with monetary wins and losses. We used different sets of stimuli in the two task runs to ensure that feedback learning was reset at the beginning of each run, and overall, 12 parallel versions of the task were used. Each version realized a specific mapping of a stimulus pair to a contingency and feedback, for the observation and control conditions respectively. The order of observation and control condition and the assignment of the different versions of the experiment to the conditions were counterbalanced across participants.

On each learning trial, participants were presented with one of three different stimulus pairs (AB, CD, EF) composed of two Japanese Hiragana characters, respectively. Participants were asked to choose one of these characters on every trial by pressing the right or left button of an MR-compatible response box. Subjects were informed that their choices determined the monetary outcomes of each trial (win or loss of 20 Eurocent) and therefore the amount of money won at the end of the experiment. Subjects did not receive explicit information regarding the probabilistic nature of the task or the exact reward contingencies. The outcome of each choice was given as symbolic feedback (either triangle or circle, counterbalanced across subjects to wins and losses (see Fig. 1). For stimulus pair AB, choosing stimulus A was associated with a win in 80% of trials, while choosing stimulus B was associated with a win in 20% of trials. Accordingly, this ratio was 70:30 for stimulus pair CD, and 60:40 for stimulus pair EF.

**Figure 1.**
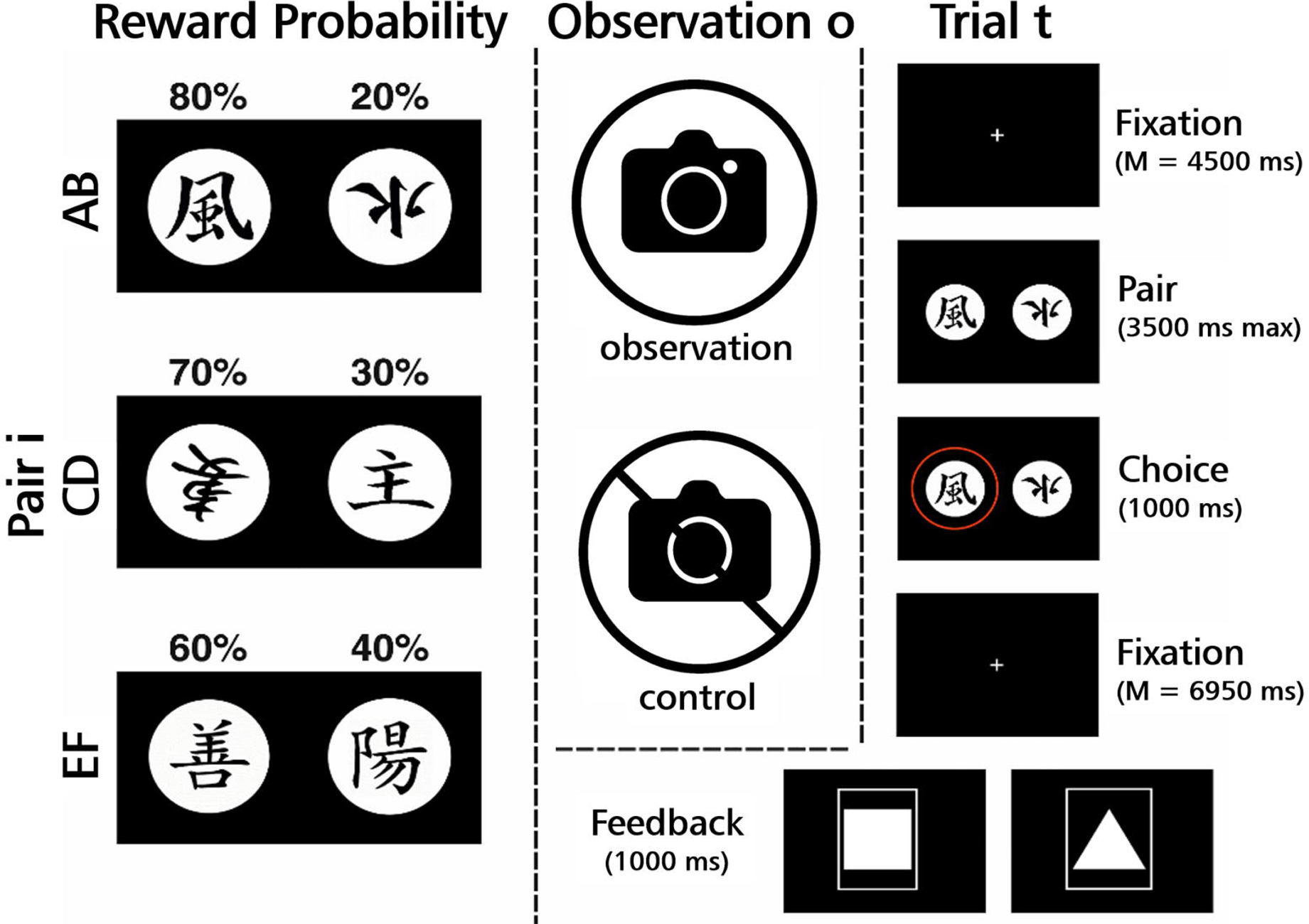

The sequence and time course of stimulus presentation in one trial of the task is depicted in Figure 1. At the beginning of each trial, a fixation cross was presented. Then, one stimulus pair (either AB, CD, or EF) was displayed until subjects made a left or right response on the response box (the response was indicated by a colored ring which appeared around the chosen stimulus of the respective pair). If response latency exceeded 3500ms, subjects were prompted to respond more quickly on future trials and the current trial was aborted. Subsequently, feedback was presented for 1000 ms. The experiment was controlled using Presentation software (Neurobehavioral Systems Inc., Berkeley, CA, USA). Each condition comprised 3 task blocks in each of which each stimulus pair was presented 20 times, thus amounting to a total of 180 trials per run and 360 trials in total.

After each of the two conditions, a *transfer of learning* (ToL) block assessed the degree to which subjects prioritized learning from wins over learning from losses or vice versa. In these TOL blocks, the symbols A (associated with a win rate of 80%) and B (associated with a win rate of 20%) were paired with each of the other stimuli C through F (associated, on average, with a win rate of 50%) to form 40 new stimulus pairs (20 involving A, 20 involving B) that had never before been encountered by the subjects. No feedback was presented during the test blocks. If participants prioritized learning from positive feedback over learning from negative feedback, they should score higher on pairs that included A than on the pairs that included B (positive transfer). If they prioritized learning from negative feedback over learning from positive feedback, they should score higher on pairs that included B than on the pairs that included A (negative transfer). The ratio *A(chosen)/B(not chosen)* therefore reflects an implicit tendency to learn from positive or negative feedback.

### Functional MRI data acquisition and preprocessing

Data acquisition was carried out with a Siemens 3 Tesla Magnetom PRISMA and a 20-channel Siemens Head Matrix Coil. Structural images were acquired using a sagittal magnetization T1-weighted (MPRAGE) sequence (TR=2130 ms, TE=2.28 ms, voxel size= 1 mm isotropic, flip angle = 8°) with 192 slices. The functional images were collected by using 210 volumes (for each of the six sessions) of a gradient-echo planar sequence sensitive to BOLD contrast (TR=2080 ms, TE=30 ms, matrix=92×92 voxel, FOV=208 mm, flip angle = 90°). Each volume consisted of 36 axial slices (thickness=3 mm, gap=0.3 mm, in-plane resolution = 2.26×2.26 mm^2^, voxel size = 2.3×2.3×3 mm). A shimming field was applied before functional imaging commenced to minimize magnetic field inhomogeneity.

All preprocessing steps were carried out using the Data Processing & Analysis of Brain Imaging (DPABI V3.1_180801, Yan et al., 2016) toolbox, which is based on SPM12 (version 7487). The first 5 data volumes were discarded due to spin saturation effects. The remaining volumes were slice-time corrected and realigned using a six-parameter (rigid body) linear transformation. The anatomical and functional images were co-registered and then segmented into gray matter (GM), white matter (WM), and cerebrospinal fluid (CSF). Functional data were spatially normalized to MNI standard space with DARTEL (Ashburner, 2007) and resampled to 2 mm isotropic voxels. Finally, the functional data were spatially smoothed with a 6 mm full width at half maximum (FWHM) Gaussian kernel.

### Models and Statistical analyses

#### Computational model of probabilistic learning

We used a standard Q-learning algorithm (Daw, 2009) to generate estimates of Q-values and RPEs from choices and the associated feedback on every trial t. For all t > 1, the model estimates trial-by-trial sequences of six expected values Q_A_, Q_B_, Q_C_, Q_D_, Q_E_ and Q_F_ for the three *stimuli pairs i* = AB, CD, EF separately in each *condition o* ∈ {observation, control}. At the start of each condition, all Q-values were initialized to 0 and trials missed due to slow responding were skipped because no gain or loss feedback was presented. For each *condition o*, a trial-by-trial update of Q of the chosen option was implemented according to the following equation:

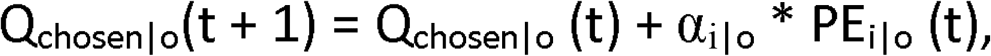

where α_i|o_ represents the learning rate of the respective pair i given the observation condition o, and PE_i|o_ represents the prediction error of the respective pair i given the observation condition o according to

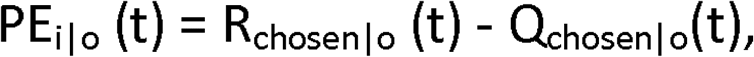

R_chosen|o_ (t) represents the presented outcome associated with the choice in that particular trial t: either +1 in case of a win or −1 in case of a loss.

The probability of the individual’s actual choice on each trial t was estimated based on the softmax rule for each pair i:

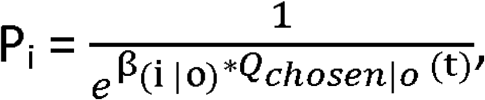

where β_(i|o)_ represents the inverse temperature controlling the degree of exploration inherent in choices.

#### General Linear Model

##### First-level single-subject analyses

Based on our hypothesis that indicators of updating during RL are altered in socially evaluative contexts in SAD patients, we set up a general linear model (GLM) with parametric modulators. This GLM accounted for 3 phases in every trial: (1) presentation of a stimulus pair, (2) participants’ choices, and (3) feedback presentation. A vector of onsets convolved with a canonical hemodynamic response function (hrf) represented each phase, respectively. Additionally, the hrf assigned to the feedback phase was parameterized by PE values stemming from RL modelling (see *Computational model of probabilistic learning*). In order to quantify differences related to observation conditions, single-subject beta-weights for these 4 regressors were estimated separately for the observation condition and the control condition, resulting in 8 task-related regressors. Additionally, we included 6 head motion parameters in the GLM.

##### Second-level random-effects group analyses

Second-level analyses were performed using permutation testing (10⍰000 permutations) in PALM (Winkler et al., 2014). First, contrasts were defined to test for main effects of RPE as well as interactions with diagnosis (SAD, HC) and interactions with condition (observation, control) and diagnosis. Based on these contrasts, t-maps were computed from subjects’ first level beta estimates, and Threshold-free Cluster Enhancement (TFCE) maps were derived as implemented in PALM. We used masks from the Brainnetome Atlas (A9m; http://atlas.brainnetome.org/bnatlas.html) and Reinforcement Learning Atlas (Pauli et al., 2018) and applied FWER-correction of p-values (α=0.05).

##### Dynamic Causal Modelling

We used Dynamic Causal Modelling (DCM) (Friston et al., 2003; Stephan et al., 2010) as implemented in SPM12 (r6591) to investigate effective connectivity. Subject-specific time series were extracted from regions of interest that were selected on the basis of the second-level random-effects (RFX) analysis. Time series were extracted from the clusters in Area 9 and VS identified by the significant interaction of RPE and diagnosis. The first eigenvariate was then computed across all voxels within these clusters. The resulting time series were adjusted for effects of no interest by specifying an F-contrast that only included the feedback condition and its parametric modulator. A bilinear one-state DCM model was constructed assuming recursive connections between both regions of interest. All feedback trials were used as driving inputs on both regions of interest. RPE parametric modulators were used as modulating inputs on the recursive connections between Area 9 and VS. DCMs were fitted to the data for observation and control conditions in both groups. To account for between-subject variability, DCMs were then subjected to a 3-level hierarchical analysis based on Parametric Empirical Bayes (PEB; Zeidman et al., 2019a, 2019b, Friston et al., 2016). The first level comprised a total of 8 DCM parameters (4 for the baseline connections (A-matrix) and 4 for the modulatory connections (B-matrix)) for every subject and observation condition which were subjected to PEB on the second and third levels, respectively. The second level comprised between-subjects effects ***X_b_***, and the third level comprised within-subjects effects ***X_w_*** (Zeidman et al., 2019b). Note that this approach accounts for differences in between-subject variability. In addition, we used a Bayesian model reduction to identify the best model given the data. The procedure derives the model evidence (free energy) by removing one or more connectivity parameters from the full PEB group-level model to produce reduced forms of the full model. With this approach, it is possible to obtain evidence for reduced models (and their respective parameters) directly from the fully-connected model and thus provides an efficient search of the model space by scoring each reduced model based on its model-evidence (for details see Friston et al., 2016). The models that maximize model evidence are then selected. After model reduction, we used Bayesian model averaging to average the connectivity parameters of the best models weighted by their evidence. That is, the most probable model will contribute the most to the average. In the result section, we report the parameter estimates of this average over the best models based on a posterior probability threshold > .99.

## Results

### Choice data

In order to analyze participants’ choices during learning, a 3 (pair: AB, CD, EF) × 3 (block: 1^st^,2^nd^,3^rd^) × 2 (condition: observation, control) × 2 (group: SAD, HC) repeated-measures analysis of variance (rmANOVA) was conducted, with Greenhouse-Geisser-correction applied whenever appropriate. The ANOVA yielded a significant main effect of pair (F[1.69, 77.78] = 7.137, p < .05) and an interaction of pair x block (F[3.33, 153.12] = 8.735, p < .05) but no other main effects or interactions (all F < 2.046). Mean percentages of correct choices according to condition and pair for SAD patients and HC are shown in Supplementary Figure 1. Within-subjects contrasts revealed significant linear trends of pair (F[1.00, 46.00] = 9.37, p < .05) as well as the interaction of pair × block (F[1.00, 46.00] = 13.13, p < .05). As no interactions with group were significant, RL from value comparisons per se seemed to be unimpaired in patients.

### Differential contributions of positive and negative feedback to learning

The TOL blocks showed (descriptive data provided in Figure 2) a 2 (feedback valence: positive, negative) × 2 (condition: observation, control) × 2 (group: SAD, HC) rmANOVA yielded a significant main effect of feedback valence (F[1.00, 46.00] = 7.959, p < .05) and a significant 3-way interaction of feedback valence, group, and condition (F[1.00, 46.00] = 6.456, p < .05). Post-hoc t-tests showed that HC learned better from negative feedback under observation (positive transfer: M(SE) = 65.10 % (± 4.90 %); negative transfer: M(SE) = 78.99 % (±3.05 %); t(23) = −2.88, p < .05), while SAD showed reduced learning from negative feedback under observation (positive transfer: M(SE) = 61.63 % (± 5.44%); negative transfer: M(SE) = 70.83 % (± 4.75%); t(23) = −2.88, p < .05).

**Figure 2.**
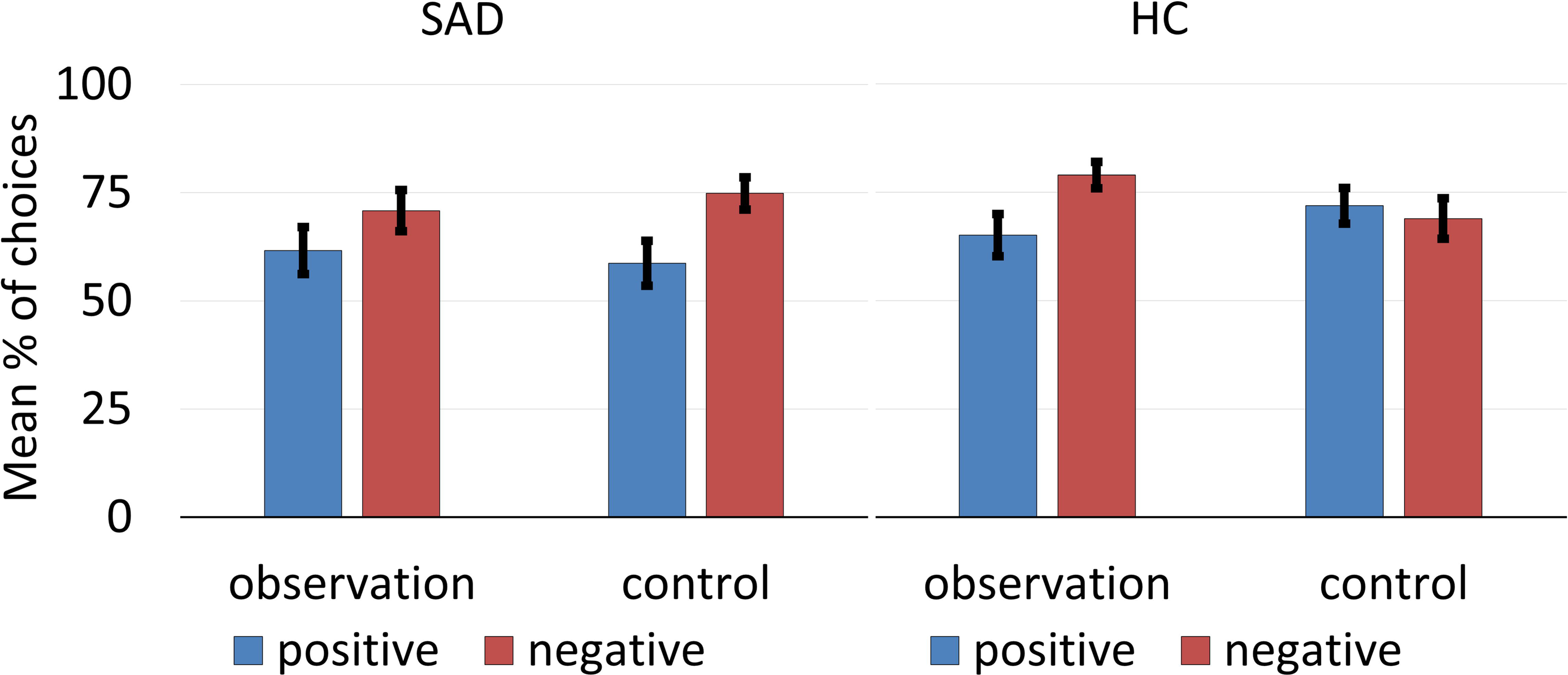

### Rating data

Under observation, SAD also reported more subjective discomfort about receiving negative feedback (M(SE) = 5.88 (± .45); t(23) = −2.64, p < .05) and higher arousal levels (M(SE) = 5.08 (± .45); t(23) = −2.40, p < .05) than HC (discomfort: M(SE) = 4.38 (± .35); arousal: M(SE) = 3.67 (± .39)).

### Learning rates

Finally, a regression analysis with learning rates onto LSAS scores was conducted (2 independent variables: (1) learning rates from the observation condition, (2) learning rates from the control condition). As expected, LSAS scores were predicted by learning rates from the observation condition (β = −.257, p_onesided_ < .05) but not by learning rates from the control condition (β = −.064, p = .67). The negative association between learning rates and symptom severity suggests slower updating of learned associations in socially anxious individuals (Figure 3).

**Figure 3.**
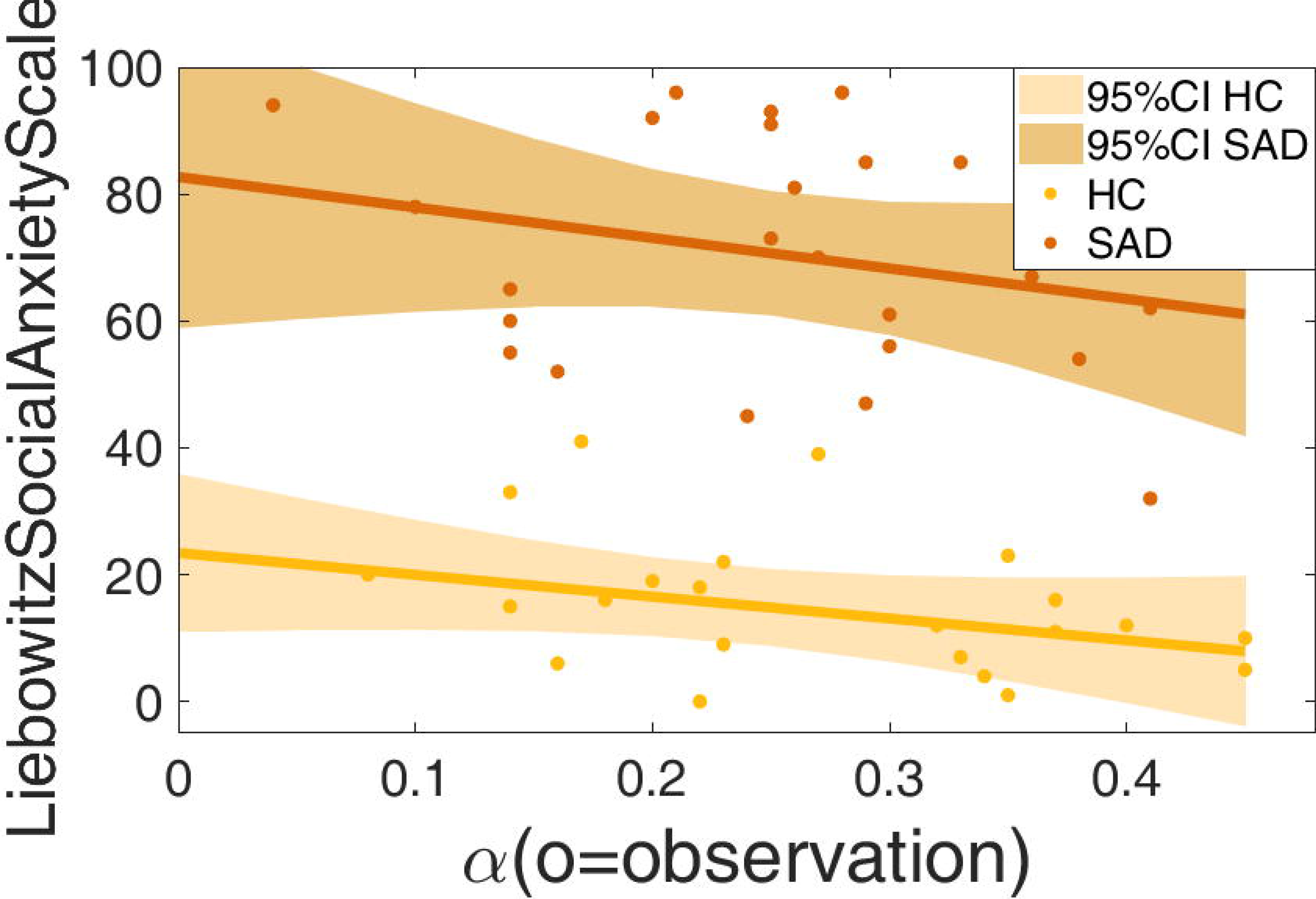

### fMRI

#### RPE-related activation

First, we tested if our model based on RPE *parametric modulators* for feedback events replicated the RL network (Pauli et al., 2018) across groups and observation conditions. A main effect of RPE-weighted feedback-related activation was found in the typical RL network comprising nucleus accumbens, lentiform nucleus (Putamen and Globus Pallidus), VMPFC, amygdala, lateral OFC, and other regions (all maps are FWE-corrected after PALM permutation analysis, p < .05). Analogous to behavioral choice data, RPE-related activation in VMPFC did not show significant effects of group or observation condition underpinning the assumption that value comparisons per se seemed to be unimpaired in SAD (see Supplementary Figure 2).

#### Differences in RPE-related activation between groups

Next, we looked for group differences in RPE-related activation. Activity of a region in DMPFC corresponding to Area 9 selectively tracked reward prediction errors in SAD (peak: x,y,z = −4,32,36). Figure 4 shows the cluster in Area 9 of DMPFC and the corresponding beta maps (averaged over all voxels within that cluster). Importantly, this same region tracked RPE in SAD preferentially when learning under observation, i.e. betas were higher in the observation than in the control condition (Fig. 5; all maps are FWE-corrected maps after PALM permutation analysis, masks used are from Reinforcement Learning Atlas and Brainnetome Atlas). Correspondingly, a subdivision of the lentiform nucleus also showed stronger coupling to RPE in SAD than in HC (peak: x,y,z = 12,8,−2; Fig. 5). Furthermore, a cluster in VS showed an interaction of RPE and group with higher activation in SAD than in HC (peak: x,y,z = −10,9,−2). The interactions in A9 and VP held after controlling for medication status as qualified by an ANCOVA with medication status as covariate (A9: F[1.00, 45.00] = .641, p = .43, VP: F[1.00, 45.00] = .003, p = .96). No other RL-relevant regions (Pauli et al., 2018) showed significant interactions of RPE, observation and diagnosis and no region showed higher activation for RPE in HC.

**Figure 4.**
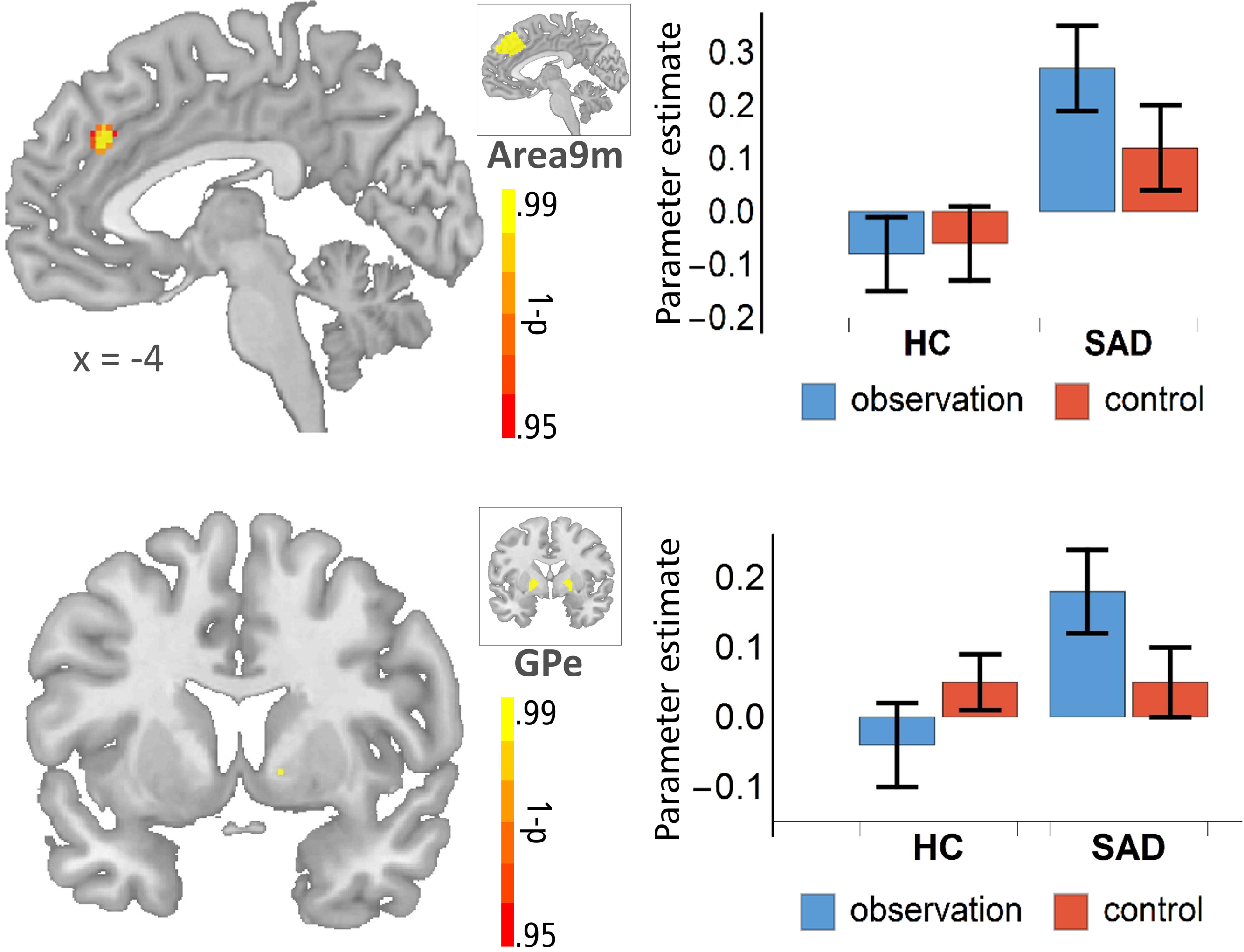

#### Session-related effects

We also asked whether learning effects modulated RPE-related activation differences between the groups. Therefore, we separately modelled predictors for each of the six learning blocks (three for the observation condition and three for the control condition). At the threshold of the other contrasts, no block-related effects were detectable.

#### Dynamic Causal Modelling

As RPEs are a likely candidate mechanism for the modulation of connection strength between regions (den Ouden et al., 2011), we used Dynamic Causal Modelling (DCM) to investigate if this mechanism showed differences between SAD and HC under observation. We specified and compared two linear models (see Figure 5): (1) we investigated if RPE signaling, comprising valence and magnitude information, alters connectivity from VS to Area 9 in SAD under observation, (2) we investigated if connectivity from VS to Area 9 was altered by positive and negative feedback (without taking into consideration the magnitude of the RPE) in SAD under observation. Model comparison was based on the Log Bayes Factor (LBF) of the difference in Free Energy of the PEB models and expressed as each model’s posterior probability. A better model fit for Model 1 (using RPE as modulatory input) over Model 2 (using only positive and negative Feedback as modulatory input) would suggest that connectivity between VS and A9 in SAD under observation is not only gated by the valence of feedback but also by magnitude information provided by our RL model. Model comparison showed strong evidence in favor of Model 1 over Model 2 (LBF = 7.75), i.e. the magnitude model best explained the data. Under observation, connectivity from VS to A9 in patients did indeed decrease as a function of RPE, i.e., positive RPEs reduced the directed connection from VS to A9 during feedback presentation while negative RPEs increased this connectivity.

**Figure 5.**
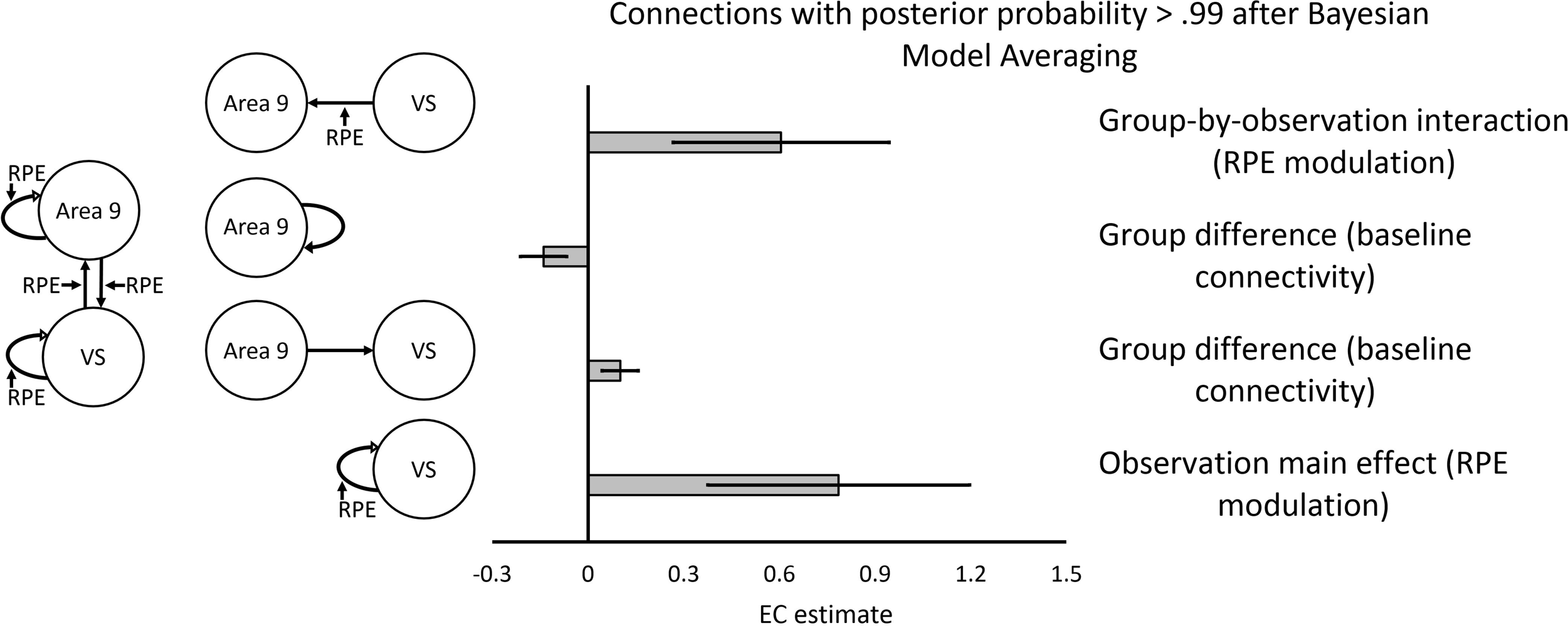

## Discussion

The present study was designed to investigate disorder-related alterations of RL in SAD brought about by social scrutiny. Therefore, patients with a DSM-IV diagnosis of SAD and HC completed a probabilistic feedback learning task (Frank et al., 2004, Voegler et al., 2019) in which they were exposed to social observation and a non-social control condition. Observation led to significantly more reported subjective discomfort in SAD than in HC, attesting to the disorder-relevance of the manipulation. Behaviorally, overall learning performance was comparable between SAD and HC, suggesting that SAD were not impaired by deficits in RL per se. However, learning rates in the observation condition predicted severity of social anxiety symptoms. Behaviorally, lower learning rates in more anxious individuals pointed towards slower updating in individuals with higher symptom burden (Browning et al., 2015, Koban et al., 2018). Assessment of TOL revealed that SAD showed a bias for negative learning under observation and in the control condition, while HC only showed a bias for negative learning under observation. This result is in line with the pattern found by Voegler et al. (2019) in an EEG study and Abraham & Hermann (2015) in a behavioral study.

On the brain functional level, RPEs during feedback presentation were expected to predict activation of a RL-network comprising ventral striatum (VS), lentiform nucleus, and VMPFC (Haber & Behrens, 2015; Pauli et al., 2018). We found comparable activation in VMPFC to RPE in both groups and both conditions. This suggests that VMPFC did not distinguish between the observation and control conditions (see also Simon et al., 2014, Becker, Simon et al., 2017). Furthermore, VMPFC activation did not distinguish between SAD and HC. Since, the percentage of correct choices was comparable in both groups across the three different contingencies AB, CD and EF, and given that VMPFC has been implicated in the coding of value comparisons, this result is in line with our previous suggestion that value computations in SAD are functionally similar to those found in control subjects. Hence, we would not assume that VMPFC contributes directly to the differences in TOL.

Based on our hypotheses, we expected SAD patients and controls to represent RPEs differently during observation in VS and Area 9 of the mentalizing network (Wittman et al., 2016, Kanske et al., 2015). In line with this prediction, we found an interaction of observation and RPE, with RPE predicting activation in this region better during observation. Finally, as expected, we found this interaction to be more pronounced in SAD, i.e., RPE in SAD during observation predicted activation in Area 9 significantly better.

The medial part of Area 9 (Fan et al., 2013, Petrides & Pandya, 1999, Sallet et al., 2013) is located in the DMPFC which is arguably the most important brain region in social inference (Amodio & Frith, 2006) and has been ascribed a particular role when choices are being made based on the alleged value standards of others in their absence (Hampton et al., 2008, Nicolle et al., 2012, see Joiner et al. for review). Area 9 has been identified as a functional subregion of DMPFC in human resting state (Li et al., 2011) and task FMRI where it is activated when confusing other’s performance with ones’s own (Wittmann et al., 2016). Our results suggest stronger coupling of Area 9 to RPEs in SAD than in HC, with this coupling further amplified by observation. As being observed increased subjective discomfort in SAD, it is interesting to ask if this discomfort was related to thinking about the expectations of the observers. Area 9 has previously been found to track expectations about the performance of others and to integrate self-related information into these expectations (Wittmann et al., 2016, see Leo & Seo, 2016 for review). This is consistent with the idea that this region is engaged when mentalizing about a dissimilar other (Mitchell et al., 2006) and learning about the social value of another player during interactions based on reliably reading others strategies (Behrens et al., 2008), imagining others’ thoughts and intentions (Saxe & Powell, 2006), or representing others’ personality traits (Van Overwalle, 2009). Nicolle et al. (2014) reported DMPFC activation not only when simulating the established value standards of another person, but also when using one’s own value standards as a proxy for the unknown value standards of others. Hence, the computational mechanisms identified by this study are in line with the cognitive mechanism central to the model of the generation and maintenance of anxiety in social/evaluative situations proposed by Rapee & Heimberg (1997). This model assumes that, in SAD, dysfunctional cognitive mechanisms during observation are characterized by biased predictions regarding the evaluation of one’s actions by others in social situations. These biases arise from dysfunctional expectations and possibly result in excessive feedback prediction error signaling. Importantly, previous studies have shown that DMPFC is a region that carries these social PEs (Behrens et al., 2008; Nicolle et al., 2012).

Prior evidence from fMRI in SAD and subclinical anxiety has implied aberrant DMPFC activation in SAD but has never shown a connection to PE signalling: Sripada et al. (2009) reported altered activation of DMPFC in SAD, albeit reduced during interactions with humans relative to a computer. In subclinical social anxiety, Peterburs et al. (2016) also found reduced activation of MPFC in HSA. However, Heitmann et al. (2014) found elevated anticipatory DMPFC activation in subclinical social anxiety to positive and negative feedback provided by human raters. Becker, Simon et al., (2017) reported a region in MPFC to show an interaction of feedback valence, observation, and SAD diagnosis. However, no study thus far has shed light on the computational mechanisms at the heart of these phenomena. The present study shows for the first time in SAD that RPEs during scrutiny can explain aberrant BOLD responses in DMPFC.

Having established this connection, the next question we asked was whether aberrant BOLD patterns in DMPFC in SAD were the source or the target of aberrant cross-talk with other regions of the RL network. VS has an important role in this network (Haber & Behrens, 2015). As VS also showed enhanced coupling to RPEs in SAD, we aimed to further elucidate the dependencies between both network regions by DCM. RPEs have been discussed as a candidate mechanism for the modulation of connection strength between regions (den Ouden et al., 2011, Stephan et al., 2010). We set up a bilinear one-state DCM using RPE as modulating input and used an hierarchical estimation procedure based on PEB to estimate its parameters. This allowed us to estimate interactions of diagnosis and observation in effective connectivity while controlling for within- and between-subject variance. The results clearly favored effective connectivity from VS to A9 modulated by RPE as the best solution over all other modulatory connections. Most importantly, the hierarchical DCM clearly showed that RPE-modulated connectivity from VS to A9 was most pronounced in SAD under observation. We tested a model containing RPE as modulatory input against a null model that only featured valence (thus disregarding the magnitude information of the RPE). This was done to ensure that the RPE was indeed a better predictor than feedback valence alone. Comparison of the FE of these models revealed that RPE magnitude significantly contributed to the explanatory power of the model. As FE punishes the number of model parameters, we made sure to keep the parameter input to both models identical in number.

Earlier studies have reported aberrant functional connectivity (FC) in SAD in DMPFC. Heitmann et al. (2016) identified hyperconnectivity of globus pallidus with DMPFC during presentation of disorder-related scenes. Gimenez et al. (2014) also reported differences in FC between SAD and HC in regions of medial frontal cortex. However, no study to date has used measures of effective connectivity to establish the directionality of aberrant regional cross-talk in SAD during probabilistic learning. Interestingly, using a very similar probabilistic learning task, Klein et al. (2007) found FC between MPFC and VS to differ for two polymorphisms of the dopamine D2 receptor gene DRD2. However, an important limitation of FC analysis is that it is based on correlational metrics and thus cannot establish the direction of connectivity (Friston, 2011). In other words, measures of FC are not able to infer true coupling, because they map consequences to causes based on statistical dependencies (Friston, 2011). For example, two regions can show substantial FC despite the absence of any true connection, just because of a common input from a third region (Friston, 2011). Moreover, since correlations depend on the level of observation noise, changes in FC arise by merely varying the signal-to-noise ratio, e.g., by increasing the number of time points or the sample size (Friston, 2011). Finally, changes in FC also arise by changing the amplitudes of neuronal fluctuations (Friston, 2011). Metrics of EC offer a way to avoid these pitfalls in network connectivity analysis. Therefore, we used DCM together with PEB to model network dependencies and their possible alteration in SAD. Other than tests based on classical statistics, PEB uses the full posterior density over the connectivity parameters from each subject’s DCM to inform results on the group-level. It thus takes into account both the expected strength of the connection and its uncertainty (posterior covariance). In other words, subjects are weighted by the precision of their estimates, such that subjects with noisy estimates contribute less to the group result. We used a hierarchical procedure with three levels to estimate the DCM parameters. We modelled baseline as well as modulatory connections between VS and MPFC on the first level, group differences between SAD and HC on the second level, and differences between the observation and control condition on the third level. This allowed us to assess interactions of diagnosis and observation in the connectivity patterns from VS to A9 at baseline and when modulated by RPE. Hence, our analysis validates the basic findings of earlier studies using FC while surmounting their main limitation in showing a clear directionality in connectivity from VS to A9.

Nonetheless, when interpreting the findings of the present study, other limitations should be kept in mind. The first limitation is the number of 24 participants in each group. However, power calculations based on published RL studies of psychiatric disorders suggest that 24 participants in each group are sufficient to estimate between-group effects of RPE in DMPFC, ACC, and VS (Cremers et al., 2012; Peterburs et al., 2016; Becker, Simon et al., 2017; Jarcho et al., 2015). Furthermore, our study has the advantage over earlier studies that we only included patients with a clinical manifestation of SAD. Hence, its validity for clinical questions can be expected to be higher. Second, it should be considered that the SAD group comprised two subjects currently treated with antidepressive medication (SSRI) as well as six subjects who received psychotherapy. While about a third of SAD patients can be expected to take SSRIs, it is not entirely clear if this might influence brain activation related to RL. However, only a relatively small number of subjects was medicated, and no direct effects of medication were statistically significant. Furthermore, a previous study using [15,2 O]-PET showed that administration of an SSRI in SAD does not generally alter the neuronal correlates of processing of social stimuli (Faria et al. 2012).

Taken together, the present study exemplifies a promising new approach to anxiety research by elucidating the role of prediction errors in SAD during social observation. A key finding is the aberrant coding of RPEs during observation in Area 9 of DMPFC – a region previously reported when using one’s own value standards as a proxy for others decisions. Current models of SAD assume this as a key mechanism in the maintenance of the disorder (Rapee & Heimberg, 1997). Hence our findings point toward altered RPE coding in Area 9 as a candidate mechanism for dysfunctional updating of expectations in SAD under observation. Moreover, we were able to show that EC between ventral pallidum and Area 9 was selectively upregulated by RPE in patients only during observation. This finding may contribute to the refinement of diagnostic criteria and differential diagnosis in anxiety disorders and might lead to a deeper understanding of the biological roots of dysfunctional expectations. Furthermore, the combination of observation manipulation by camera and computational modelling might offer a new approach for theranostics, which uses the decoupling of PEs and DMPFC activation under observation as a marker for therapeutic progress.

## Acknowledgments and Disclosures

A first draft of this manuscript has been published as a preprint on BioRxiv:

The authors report no biomedical financial interests or potential conflicts of interest.

