## Supplemental Materials for "Reward Prediction Error Signaling during Reinforcement Learning in Social Anxiety Disorder is altered by Social Observation"

Supplementary Material

### Supplementary Figure 1

*Mean percentage of correct choices during learning according to condition (observation/control) and pair (AB, CD, EF) for HC (left) and SAD (right). Error bars represent standard errors.*

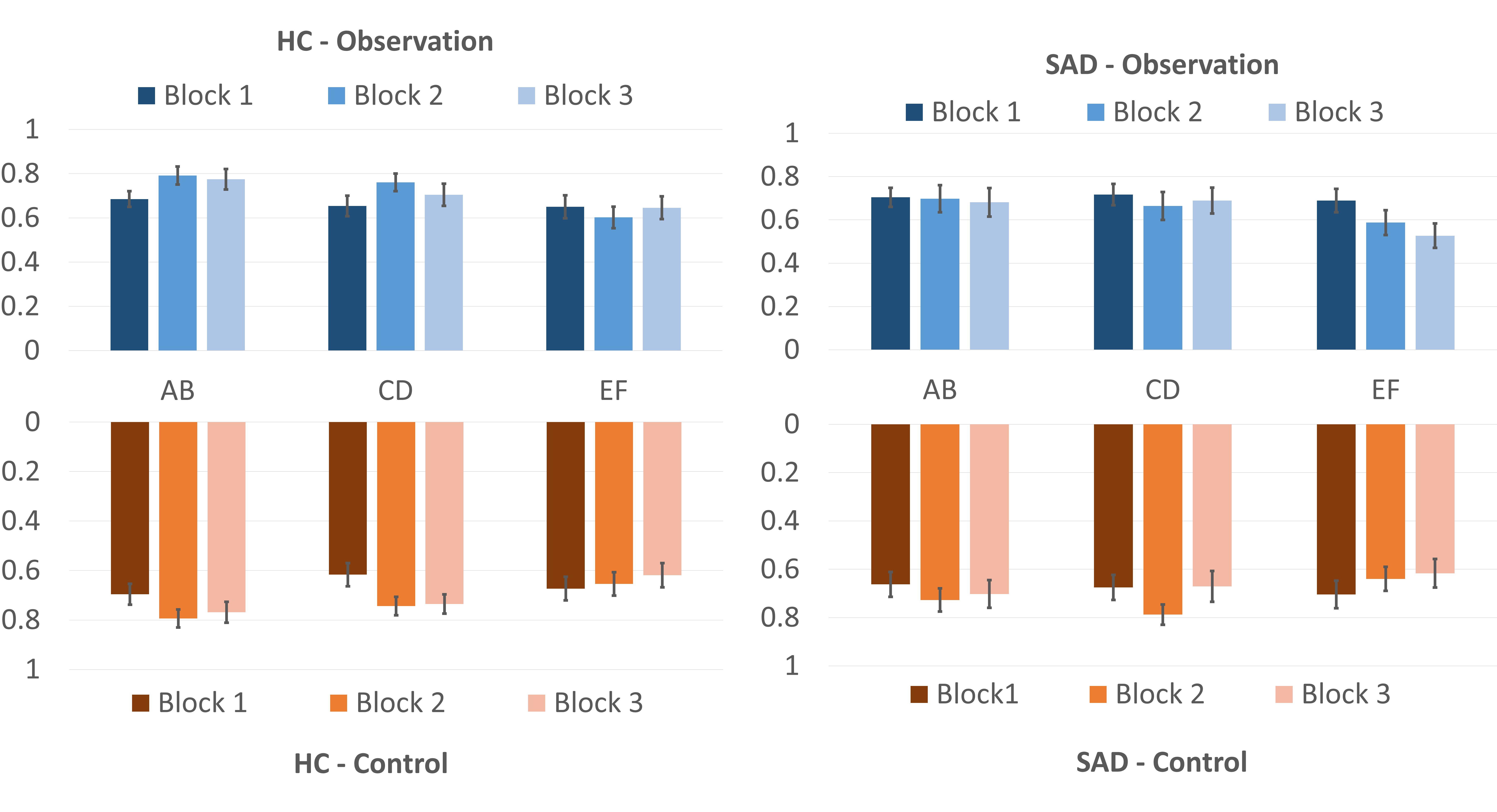

### Supplementary Figure 2

*Main effect of RPE in VMPFC.* Below: *Beta plots for observation and control conditions in SAD and HC. Error bars are median standard errors (SEs).*

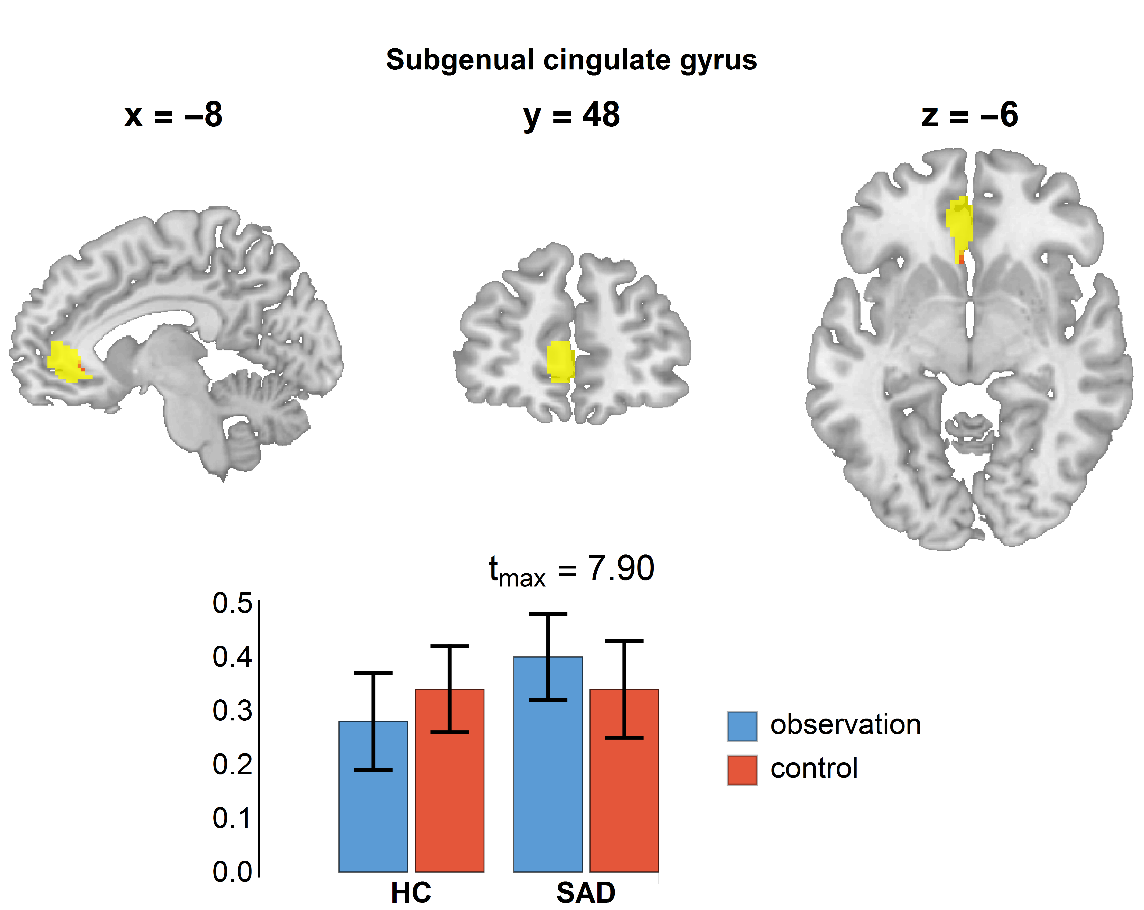

### Supplementary Figure 3

Multilevel PEB Design matrix and corresponding regressors: The first regressor models the mean of the group means, the second the mean of treatment effects related to observation and control condition, the third regressors models the difference between group means and the last regressors models the group difference of the difference between observation and control condition.

| **Regressors** | **Model design matrix** | | | |
| --- | --- | --- | --- | --- |
| Mean (Mean(*SAD*), Mean(*HC*)) | 1 | 0 | 1 | 0 |
| Mean (*SAD_(observation - control)_, HC_(observation - control)_*) | 0 | 1 | 0 | 1 |
| Mean(*SAD*) - Mean(*HC*) | 1 | 0 | -1 | 0 |
| [*SAD_(observation - control)_*] - [*HC_(observation - control)_*] | 0 | 1 | 0 | -1 |

### Supplementary Figure 4

Output of Bayesian model reduction. **Top row**: log-posterior (left) and posterior probability (right) of the best 235 models. **Middle row**: Maximum a posterior (MAP) estimates of the group-level PEB parameters before (left) and after (right) the search through the model space. **Bottom row**: model space of the best 235 models (left) and posterior probability of each parameter (right). The probability is obtained by comparing the evidence for all models (from the best 235) with this parameter switched on, versus all models with this parameter switched off.

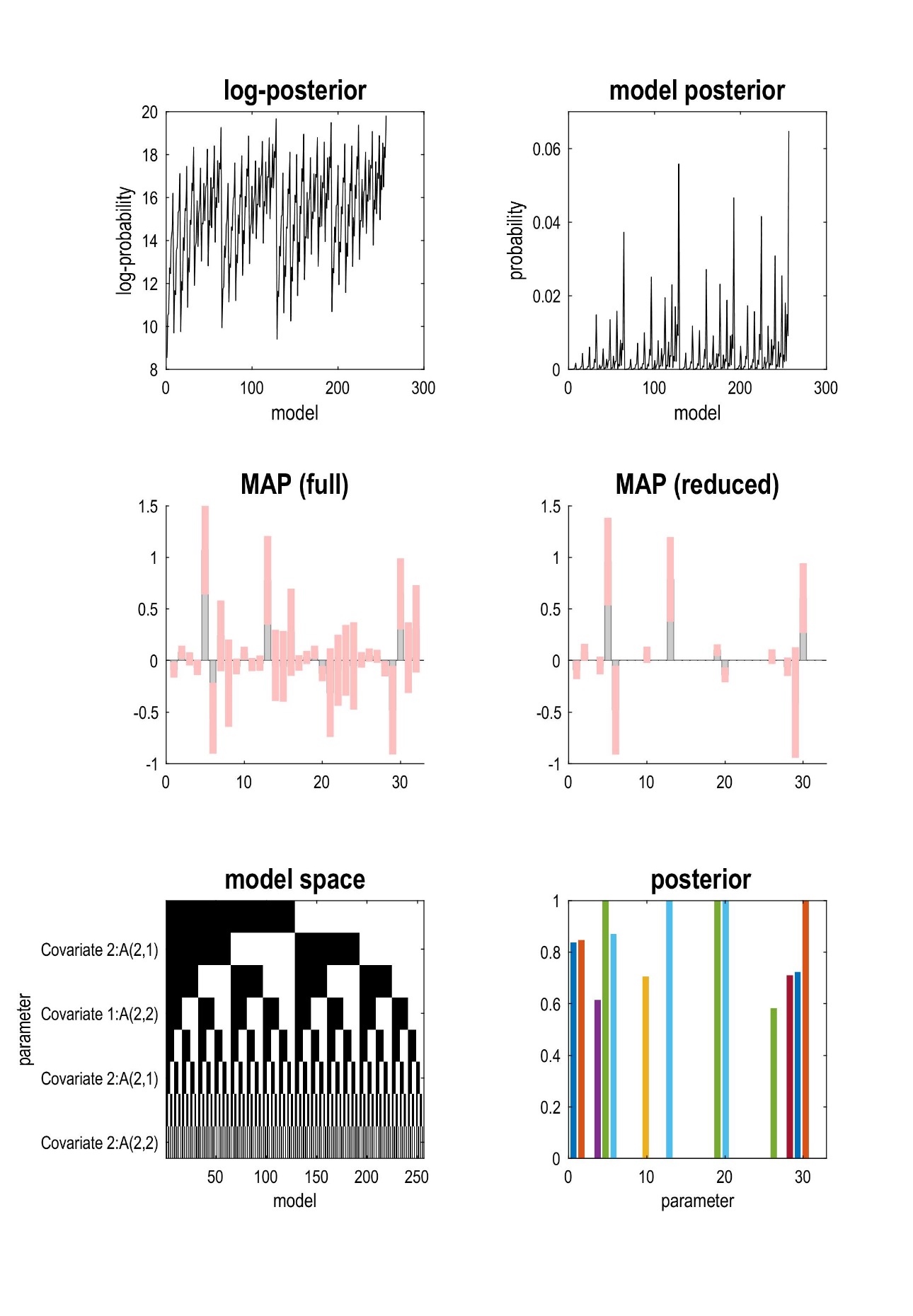

### Supplementary Figure 5

Output of Bayesian model comparison. **Top row**: Effect sizes of the EC parameters for Covariate 1 (left), which represents commonalities across groups, i.e. mean of group means (A-matrix is shown in parameter 1 to 4, B-matrix in parameter 5 to 8 and mean of treatment effects (observation – control) over both groups, A-matrix is shown in parameter 9 to 12, B-matrix in parameter 13 to 16) and Covariate 2 (right), which represents differences between groups, i.e. difference between group means (A-matrix is shown in parameter 1 to 4, B-matrix in parameter 5 to 8) and difference in treatment effects between groups (A-matrix is shown in parameter 9 to 12, B-matrix in parameter, 13 to 16) before model reduction. **Middle row**: Effect sizes after reducing the parameters to the most relevant, i.e., that contributed most to model evidence for the parameter estimates of Covariate 1 (left) and Covariate 2 (right). **Bottom row**: Posterior probabilities for the parameters of Covariate 1 (left) and Covariate 2 (right).

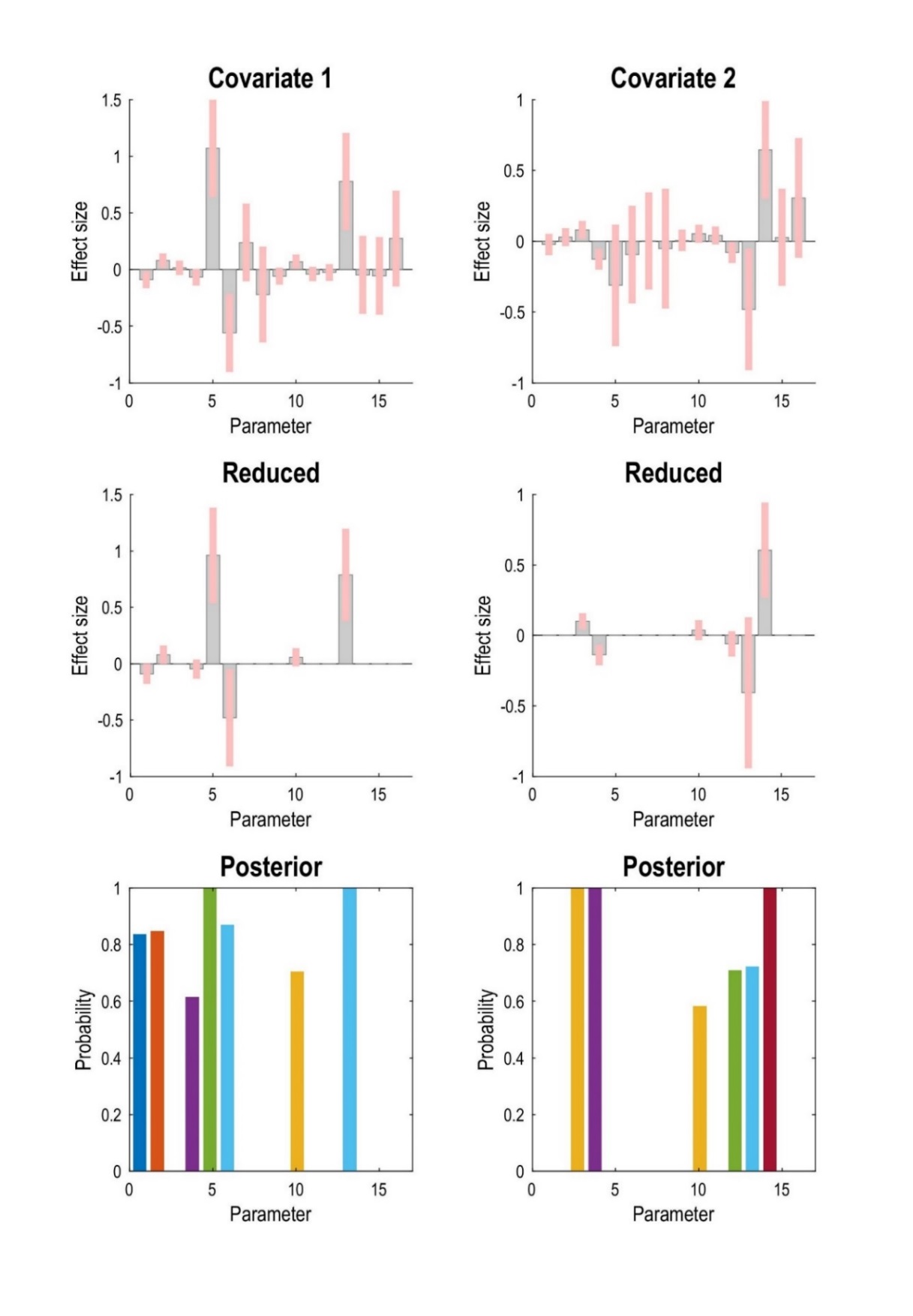
